# Reduced transfer of visuomotor adaptation is associated with aberrant sense of agency in Schizophrenia

**DOI:** 10.1101/338509

**Authors:** Sonia Bansal, Karthik G Murthy, Justin Fitzgerald, Barbara L. Schwartz, Wilsaan M. Joiner

## Abstract

One deficit associated with schizophrenia (SZ) is the reduced ability to distinguish sensations resulting from self-caused actions from those due to external sources. This reduced sense of agency (SoA, awareness of ownership over self-generated actions) is hypothesized to result from a diminished utilization of internal monitoring signals of self-movement (i.e., efferent copy) which subsequently impairs forming and utilizing sensory prediction errors (differences between the predicted and actual sensory consequences resulting from movement). Here, we investigated the connections between clinical SZ symptoms and motor adaptation, a process that uses sensory prediction errors to update motor output. Schizophrenia patients (SZP, N=30) and non-psychiatric healthy control subjects (HC, N=31) adapted to altered movement visual feedback, and then applied the motor recalibration to untested contexts (i.e., the spatial generalization to untrained targets). Although adaptation was similar for SZP and controls, the extent of generalization was significantly less for SZP; movement trajectories made by patients to the furthest untrained target (135°) before and after adaptation were largely indistinguishable. Interestingly, deficits in the generalization were correlated to positive symptoms of psychosis (e.g., hallucinations), but not negative symptoms. Generalization was also associated with subjective measures of SoA across both SZP and HC, emphasizing the major role action awareness plays in motor behavior, and suggesting that tendencies to misattribute agency, even in HC, manifest in abnormal motor performance. We discuss the possible link of these findings to cerebellar circuit abnormalities that may be a common source for deficits in the utilization of sensory prediction errors and aberrant SoA.

## INTRODUCTION

Adaptation paradigms have been used to investigate how motor behavior is modified in response to misalignments between intended body motions and subsequent sensory consequences. As a result, various properties of the motor recalibration process have been demonstrated, such as the spatial generalization (i.e., transfer of learning to different movement directions, Berniker *et al*. 2014; Brayanov *et al*. 2012; Fernandes *et al*. 2012, 2014; Taylor *et al*. 2013; Zhou *et al*. 2017). These generalization patterns provide insight into the internal representations of the learning (Poggio and Bizzi, 2004; Shadmehr, 2004). For instance, the spatial transfer generally decreases exponentially with the distance away from the trained direction, suggesting the underlying neural basis functions are rather narrow (Brayanov *et al*. 2012; Zhou *et al*. 2017).

Based on these studies, adaptation features, such as spatial generalization, may provide insight into the properties and possible deficits in the neural control of movement and different motor learning mechanisms. Schizophrenia (SZ) is a neuropsychiatric disorder associated with defects in utilizing internal signals (e.g., the efferent copy of issued motor commands) to predict the result of action prior to (or in the absence of) sensory feedback. Impairments in these signals have been implicated in the clinical symptoms of SZ, specifically difficulty in distinguishing internal from externally-derived experiences and disturbed sense of agency (SoA, i.e., ownership of self-generated actions and their outcomes) (Blakemore *et al*. 2002; Feinberg, 1978; Ford and Mathalon, 2005; Frith *et al*. 2000). Correspondingly, schizophrenia patients (SZP) have difficulty in tasks that depend on internal predictions (Bansal *et al*. 2018b; Daprati *et al*. 1997; Lindner *et al*. 2005; Martinelli *et al*. 2017; Shergill *et al*. 2005, 2014). For instance, Synofzik *et al*. (2010) tested subjects in a task in which external visual feedback was rotated with respect to the actual unseen movement. Following the movement, subjects were asked to report the rotation direction in relation to the movement, providing a detection threshold. Normal detection depends on integrating two information sources: the internal prediction and external sensory feedback (Shadmehr *et al*. 2010; Synofzik *et al*. 2013). Consistent with an internal prediction deficit, SZP were less accurate than healthy controls (HC), exhibiting higher detection thresholds that correlated with positive SZ symptoms (e.g., delusions). Thus, in SZ, abnormal internal predictions of movement outcomes may result in a reduced sense of agency over actions, and a subsequent difficulty in properly detecting and adjusting to movement errors—especially when these adjustments are tested in untrained/unfamiliar contexts.

Here, similar to Synofzik and colleagues (2010, 2013), to assess the role of action ownership in motor learning and generalziation, we utilized a visuomotor rotation paradigm in which subjects adaptated arm reaching movements to rotated visual feedback. Specifically, we investigated the spatial (the adaptation transfer to different movement directions) and temporal properties (the adaptation transfer over successive trials) of generalization. Consistent with previous studies (Bigelow *et al*. 2006; Ninomiya *et al*. 1998; Rowland *et al*. 2008), there was no significant difference between SZP and HC in the rate of adaptation. However, compared to HC, SZP were impaired in the adaptation transfer to different movement directions; unlike HC, movements to the furthest target (135°) before and after adaptation were not significantly different for SZP. In addition, deficits observed in the spatial generalization were correlated to positive (but not negative) symptoms of SZ and disturbed SoA. Interestingly, the extent of motor adaptation transfer was also associated with subjective measures of SoA in HC, suggesting that an accurate sense of action ownership is an important component in applying the motor recalibration to untested contexts. We discuss how this behavioral impairment may involve cerebellar circuit abnormalities that possibly affect both the sense of action ownership and error-based motor learning.

## MATERIALS AND METHODS

### Participants

Thirty-one veterans who met Diagnostic and Statistical Manual of Mental Disorders, Fourth Edition DSM-IV criteria for schizophrenia or schizoaffective disorder (SZ) were recruited from the Washington DC Veterans Affairs Medical Center (DCVAMC). One patient was unable to operate the equipment during testing and was thus excluded. Subjects were outpatients receiving stable doses of either typical (N = 1) or atypical (N = 28) antipsychotic medication, or an antidepressant (N = 1) for at least three months prior to testing. Chlorpromazine (CPZ) equivalent antipsychotic dosages were calculated for each patient (Woods, 2003). Thirty-one healthy, control participants (HC) with no psychiatric condition and no personal or family history of mental illness diagnosis were recruited from the DCVAMC and enrolled in the control group. All participants were screened to exclude substance use, neurological disorders and history of head injury. All participants had normal or corrected-to-normal vision. Groups were matched for age, sex, and handedness. The Institutional Review Boards (IRB) at the DCVAMC and George Mason University approved this study and written informed consent was obtained from all participants prior to testing. Participants were reimbursed for participation. Demographic information is presented in Table 1.

**Table 1.** Summary of participant demographics.

|  | HC, N=31 | SZP, N=30 |  |  |
| --- | --- | --- | --- | --- |
|  | Mean (SD) | Mean (SD) | Statistic | P |
| Age (yrs) | 54.81(9.01) | 55.76 (9.84) | $t = 0.06$ | 0.95 |
| Gender | 3 F / 28 M | 2 F / 28 M | $\phi = 0.18$ | 0.66 |
| Handedness | 3 L / 28 R | 6 L / 24 R | $\phi = 1.29$ | 0.26 |
| Years of education | 14.89 (2.33) | 13.69 (2.58) | $t = 1.89$ | 0.06 |
| Standard WRAT Scores<br>(Estimation of IQ) | 100.19 (8.82) | 94.57(13.13) | $t = 1.97$ | 0.05 |
| Total SoA Score | 33.94 (1.15) | 39.60 (7.97) | $t=3.91$ | <0.005 |
| “Physical” SoA | 9.03 (0.41) | 12.32 (3.88) | $t=4.69$ | <0.001 |
| “Mental” SoA | 13.74 (0.85) | 17.63 (5.92) | $t=3.81$ | <0.005 |
| “Social” SoA | 11.16 (0.33) | 9.66 (2.76) | $t=3.05$ | <0.005 |
| Duration of illness (yrs) |  | 28.30(9.94) |  |  |
| CPZ Equivalent Dose<br>(mg/day) |  | 462.42 (261.57) |  |  |
| PANSS Total |  | 57.54 (10.04) |  |  |
| PANSS Positive |  | 16.35 (5.85) |  |  |
| PANSS Negative |  | 16.57 (5.66) |  |  |
| PANSS General |  | 26.09 (6.14) |  |  |

Data are the mean ± SD.
SoA, Sense of Agency; WRAT, Wide Range Achievement Test; PANSS, Positive and Negative Symptoms Scale; CPZ, chlorpromazine equivalent of total antipsychotics.

### Clinical Measures

Clinical symptoms in the patient group were assessed using the Positive and Negative Syndrome Scale (PANSS; Kay *et al*. 1987), which consists of three main subscales: Positive Symptoms, Negative Symptoms and General Symptoms. In addition, the Sense of Agency Scale (SOAS; Asai *et al*. 2009a, b) was used to assess sense of agency in both patients and controls (Bansal *et al*. 2018a) and consists of three subscales: misattribution of the agent (mental agency), uncontrollability of one’s own body (physical agency), and assertiveness in social situations (social agency). The Wide Range Achievement Test (WRAT) (Wilkinson and Robertson, 2006) was used to estimate premorbid IQ.

### Experimental Setup

The experimental setup was similar to that used in Wu and Smith (2013) and Zhou and colleagues (2017). Subjects were seated at a desk in an adjustable chair facing a horizontal LCD monitor (BenQ XL2720T). The chair height was adjusted for each subject to ensure that they could comfortably see the monitor and reach towards all presented target positions. The experimental system included the monitor, a digitizing tablet and a PC to run the experimental paradigm and collect the behavioral data. The LCD monitor was mounted horizontally in front of subjects at shoulder level, displaying the various visual cues during the experiment. The monitor was 8 inches above the digitizing tablet (a workspace of 12 inches by 19 inches; Intuos3, WaCom) that was used to track and record hand position at 200 Hz. The monitor and tablet were held by an aluminum framing system (80/20 Inc.) with outer dimensions of approximately 26 × 18 inches. Subjects grasped a cylindrical handle (2.5 cm in diameter) that was embedded with a stylus, and made reaching movements on the tablet. The set-up ensured that subjects only viewed the monitor screen during the movements; the position of the monitor obstructed the view of the tablet. The handle had a flat bottom covered with Teflon, allowing it to glide smoothly on the tablet with little friction. The stylus/hand position was represented as a screen cursor (2.5 mm in diameter). The midline of the participant was approximately aligned with the center of the tablet and monitor. This also served as the center of the workspace.

### Intralimb spatial generalization of visuomotor adaptation

The paradigm was similar to the task structure in Zhou *et al*. (2017). Starting from an initial start target (5 mm in diameter) located in the center of the workspace, subjects moved the cursor 9 cm to a reach target (7 mm in diameter) also located along the midline. Subjects were instructed to make direct movements towards the target as though they are reaching for an object. Subjects received visual and auditory feedback on movement speed. If the movement was too slow (≥ 700 ms in duration) the target turned blue once the radial distance was exceeded. If the movement was too fast (≤ 250 ms) the target turned red. The target turned green for movement durations that were between 250 and 700 ms. In addition, a short duration beep (at a frequency of 429 Hz) was given to signify a good trial.

Subjects first completed an initial *familiarization phase*. Subjects made reaching movements to peripheral targets located at a radial distance of 9 cm at seven target locations (0°, ±15°, ±75° and ±135°) (Figure 1C). This block of trials consisted of 28 movements with full visual feedback, 4 movements to each of the 7 target locations. During these trials the cursor followed the true position of the hand. Following the familiarization phase, subjects completed the *baseline phase*, which consisted of 70 trials split into 2 blocks. During each block, movements to each target were repeated 5 times, with visual feedback removed for 3 of the 5 movements. These trials with no visual feedback (6 for each of the 7 target locations over both blocks), served as the baseline for the generalization probe trials described below.

**Figure 1.**
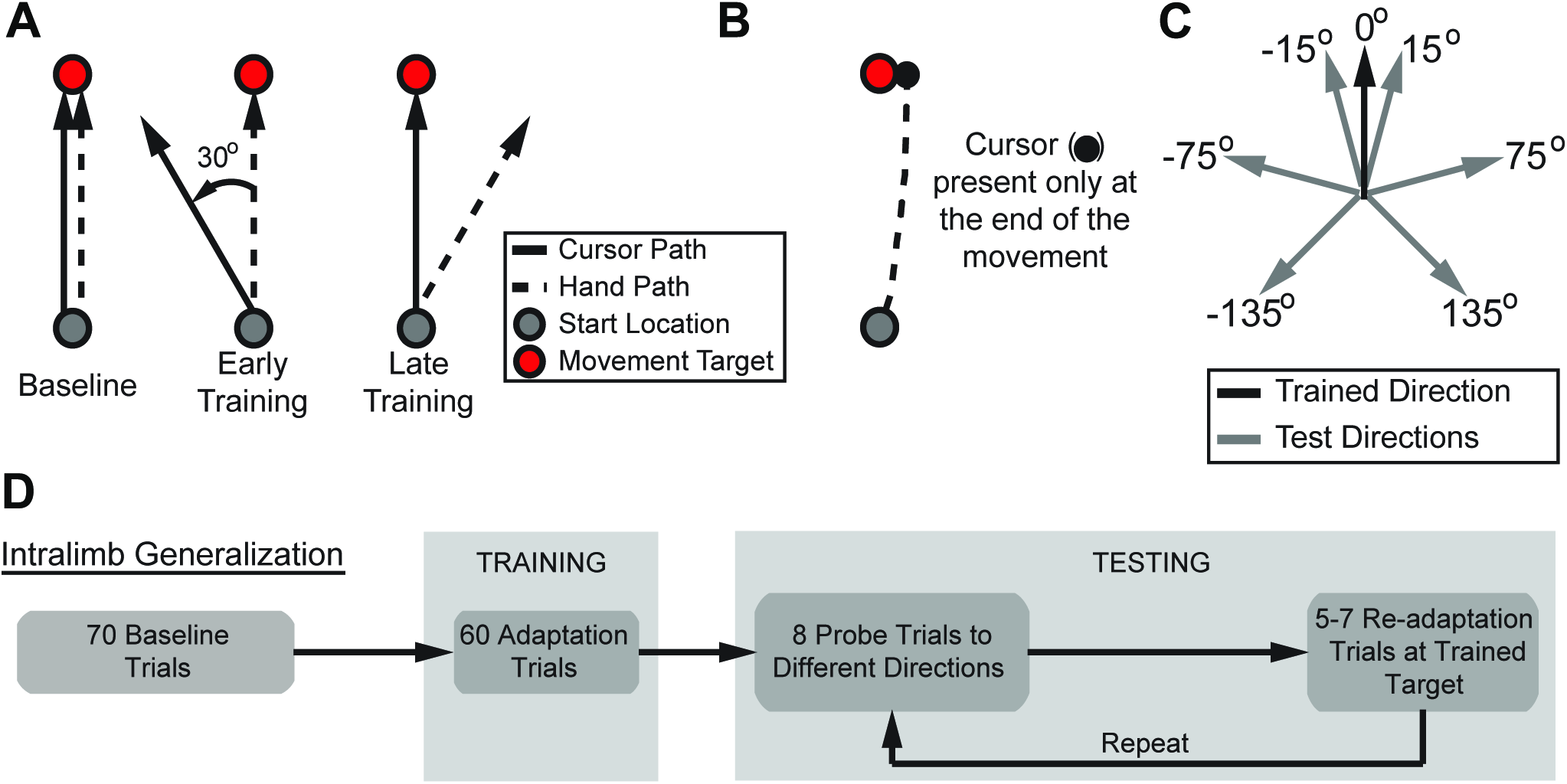
Experimental paradigm. (Figure adapted from Zhou *et al*. 2017). **(A)** Representation of the hand and cursor movements before, during and at the end of training. During the baseline phase the cursor and hand position are aligned. Once the perturbation is applied during the training phase there is a rotational offset between the cursor and hand movement. Late in training there is a compensatory rotation in the hand trajectory that is applied in order to direct the cursor to the target. **(B)** When the perturbation is applied endpoint visual feedback for the movement is provided once the movement exceeds the target distance of 9 cm. **(C)** Movement directions to test spatial generalization. Participants were tested on peripheral targets located at a radial distance of 9 cm at seven locations (0° (Trained direction), ±15°, ±75° and ±135°). **(D)** Experimental sequence. There was an initial baseline phase, followed by a visuomotor rotation training phase. Following training there was a testing phase in which the spatiotemporal properties of the trained visuomotor adaptation were assessed.

After the baseline phase, subjects completed the *training phase* during which a visuomotor rotation (VMR) was applied to the cursor feedback. On these trials the cursor path was rotated around the hand path by either *θ* = +30° (clockwise, CW) or *θ* = ‒30° (counterclockwise, CCW) (Figure 1A). The first 15 trials of the block were completed without the cursor rotation; the perturbation was applied abruptly on the 16^th^ trial for 60 total trials with the perturbation. Below we will refer to these 60 perturbation trials when describing performance during the training. On trials with the rotated visual feedback subjects were only provided endpoint feedback; the cursor was extinguished at movement initiation (when hand velocity exceeded 5 cm/s) and reappeared along an invisible circle once the movement exceeded the radial distance of the target (9 cm) (Figure 1B). This feedback was presented for 1.5 seconds and then was replaced by a circle centered on the initial start target. The radius of the circle matched the distance of the hand from the start position. Subjects moved their hand back to the start position guided by the size of the circle; the circle radius decreased in proportion to the distance from the start target. This limited the spatial information of the actual movement trajectory, but still provided enough information for the subjects to return to the start target. Once the hand was within 1 cm of the start position, the cursor reappeared and subjects were instructed to place the cursor in the start target for one second to begin the next trial.

After training, subjects completed the *testing phase* during which we assessed generalization of adaptation to the seven different movement directions, spanning ±135° around the trained movement direction (Figure 1C). First, subjects made 8 movements to (pseudo randomly selected) assess transfer (generalization probe movements). There were 7 possible targets (0°, ±15°, ±75° and ±135°) ensuring that at least one movement direction was tested twice during these generalization probe movements. We structured the experiment so that at the end of the session each of the 7 movement directions was experienced at each of the 8 different placements within the probe sequence. That is, each movement direction was tested for generalization early (first probe), late (eighth probe) and all the times in between (two through seven). During these probe movements subjects received no visual feedback on performance (blank trials). These 8 generalization probe movements were followed by 5-7 retraining trials during which subjects made movements to the trained target with the applied visual rotation and were provided endpoint visual feedback. This pattern (5-7 retraining movements followed by 8 generalization probe movements, Figure 1D) was repeated throughout the session, for a total of 8 generalization probe movements for each of the 7 targets. Due to the randomness of the retraining trials, subjects made a total of between 96 to 112 movements that were divided between two blocks.

### Analysis

We determined the direction of each movement by calculating the angle of the line connecting the hand position at the start of the movement and 4.5 cm into the reach, and the line from the center of the start location to the center of the peripheral target. Angular bias based on the average of the baseline trials was first subtracted from these adaptation and generalization probe movements. These movement angles during training were compared for early, middle and late adaptation by averaging a bin of 3 trials at the start (early, trials 1-3), middle (middle, trials 30-33) and end (late, trials 58-60) of the visuomotor rotation training phase (Figure 2A). The transfer of adaptation on generalization probe trials was quantified by the ratio of the (baseline subtracted) angular deviation of movements made on the blank generalization probe trials to the average angular deviation (baseline subtracted) on the preceding retraining trials, specifically the last 3 trials of the 5-7 retraining sequence. This ratio was then scaled to provide a percentage of adaptation transfer. Note that by computing this percentage, the transfer of adaptation is relative to the angular deviation during retraining. Thus, the percentage is computed the same for both CW and CCW rotations. The spatial change of adaptation transfer (Figure 3A) was determined as the change in percent adaptation transfer at the trained target as a function of absolute angular distance away from trained target (Figure 3B). Here we scaled the percentage of transfer amount by the mean percent transfer at the trained target location. The temporal decay of adaptation at the trained location (Figure 5) was determined as the angular deviation at the trained target location over generalization probe order (1 through 8). Similar to the spatial changes in adaptation transfer, we normalized the amount by the group mean percent transfer on the first probe trial. Except where noted, we provide the mean across subjects and the standard error of the mean.

**Figure 2.**
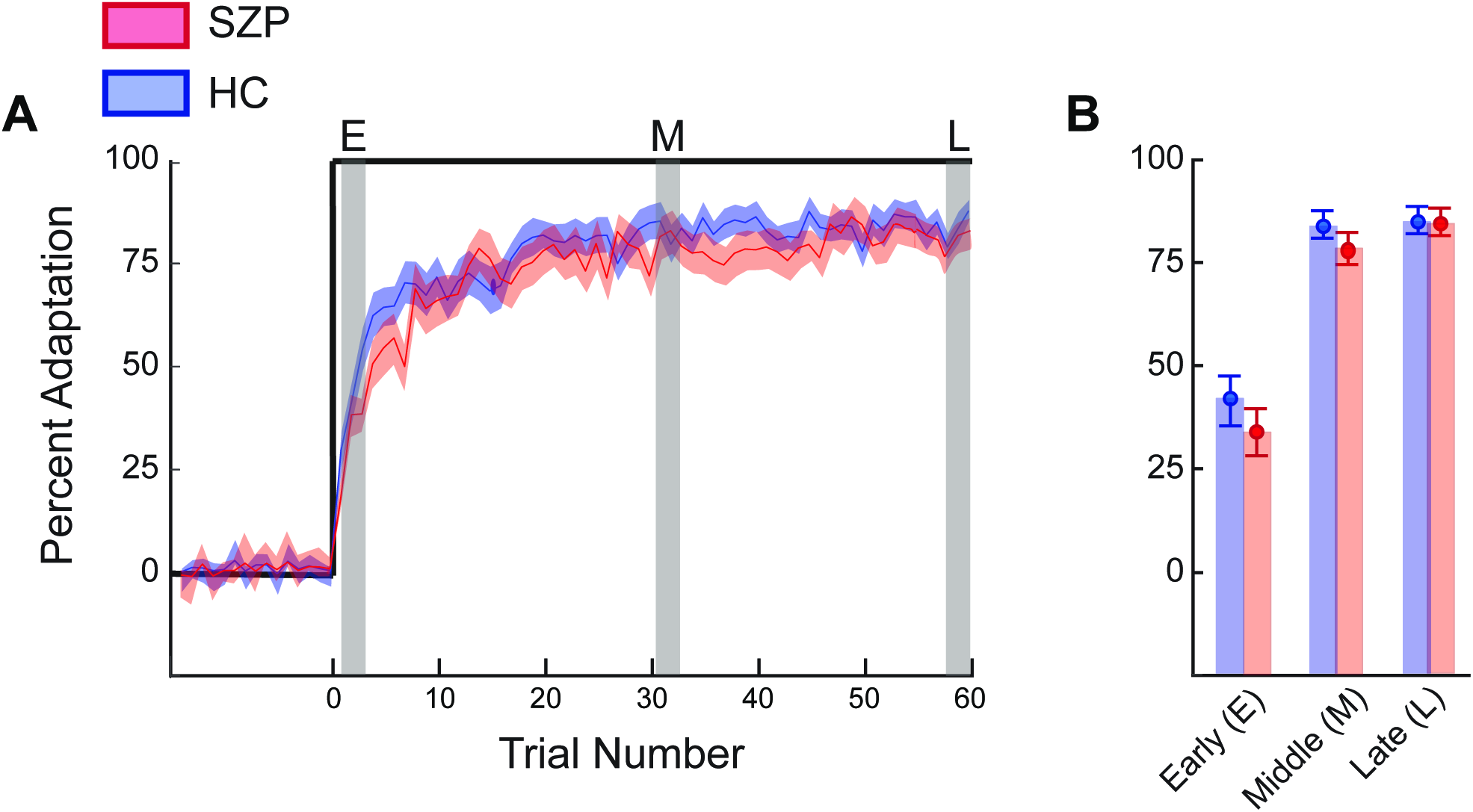
Adaptation to visuomotor rotation during training. **(A)** Adaptation curves over the training phase. Both SZP and HC (red and blue traces, respectively) were able to adapt to the 30° visuomotor rotation by adjusting the reach angle by approximately 26° by the end of the training phase (percent adaptation of approximately 86%). The solid lines represent the mean percent adaptation and shaded regions are the standard error. **(B)** Adaptation level during early, middle and late periods of visuomotor training. The percent adaptation for the two groups were compared for early, middle and late adaptation by averaging a bin of 3 trials at the start (trials 13), middle (trials 30-33) and end (trials 58-60) of the training phase. The vertical bars represent standard error. Across all three periods there was no significant difference in the adaptation levels between groups.

**Figure 3.**
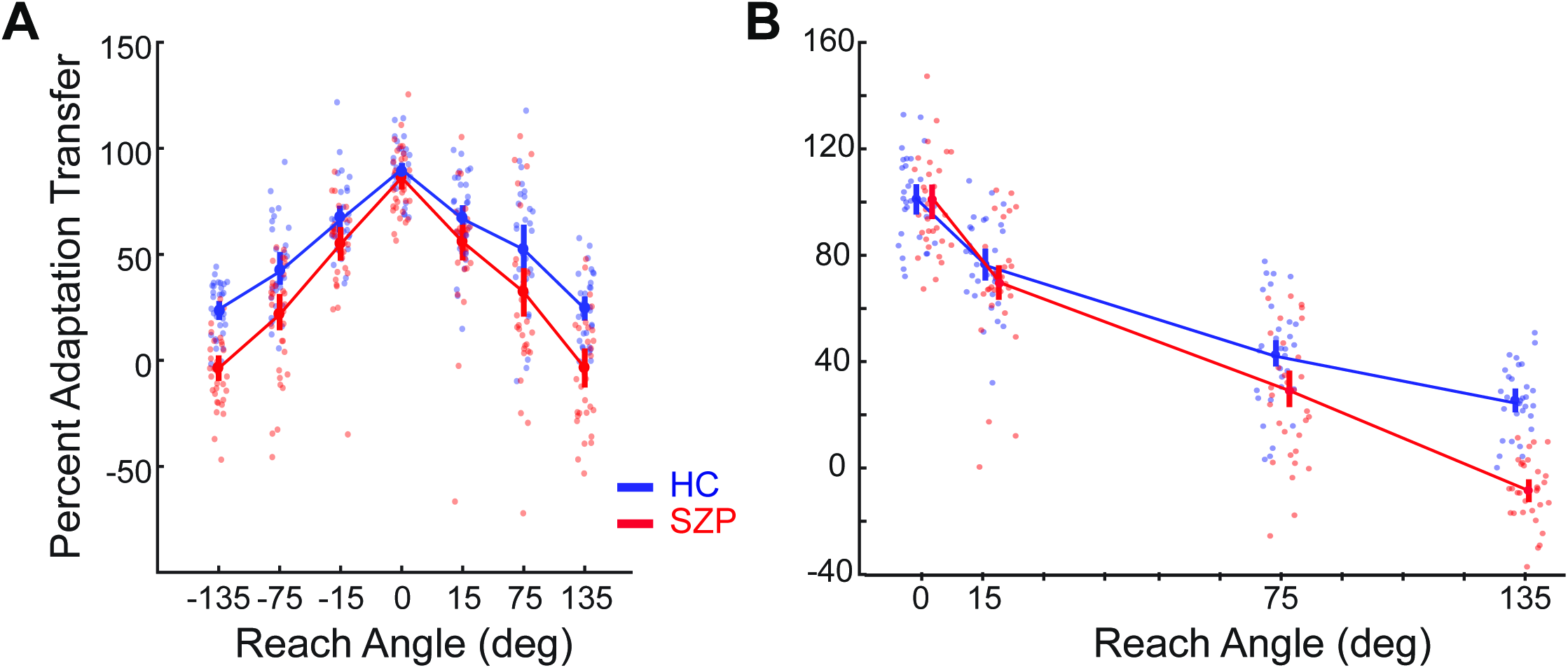
Spatial generalization of visuomotor adaptation. **(A)** The spatial generalization (percentage of adaptation transfer) to each of the targets for both groups (HC are represented by blue symbols, SZP are red). The percentage of adaptation transfer was determined on the generalization probe trials relative to the prior retraining trials. Each small filled circle represents the results of a single subject, larger filled circles represent the mean level for each respective group and vertical lines represent standard error. **(B)** Normalized spatial generalization. The generalization data in panel A were combined across workspace locations (absolute angular distance away from trained target) and normalized so that at the trained direction (0° target location), the adaptation is 100% by dividing each subject’s transfer percentage by the group mean. The adaptation to targets at 15°, 75° and 135° away from the trained direction were scaled relative to this normalized percent value. As in panel A, each filled circle represents the results for a single subject, larger filled circles represent the mean level for each respective group and vertical lines represent standard error.

**Figure 4.**
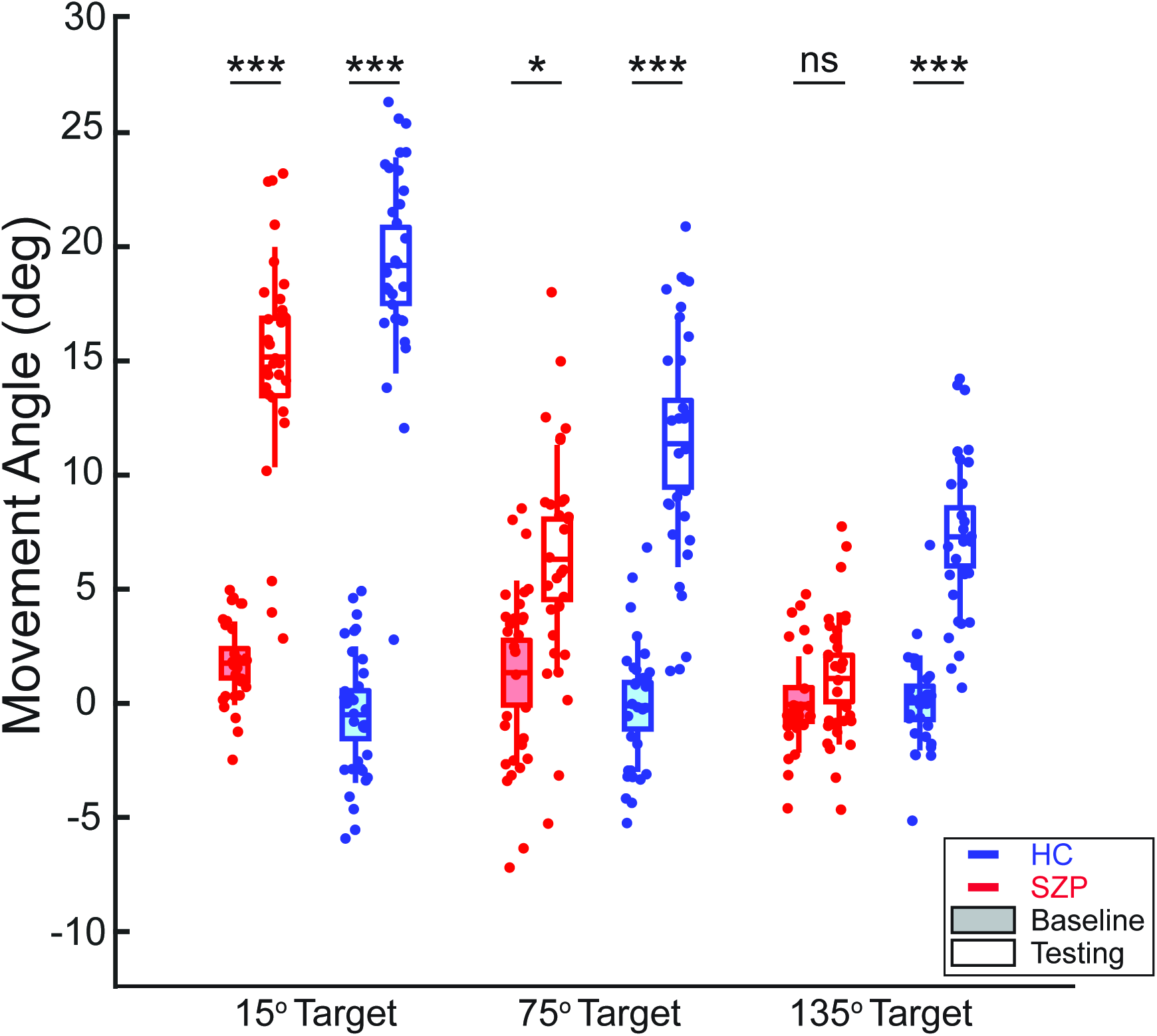
Comparison between baseline and post-adaptation testing movements. We examined differences in movement direction between baseline and post-adaptation testing trials in both groups (SZP depicted by the red symbols, HC by the blue symbols) for targets away from the trained location (15°, 75° and 135°). Each filled circle represents the results of a single subject. Filled boxplots indicate mean movement angles at baseline (full visual feedback, before visuomotor rotation training), and unfilled boxplots represent the mean movement angles in the testing phase, with no visual feedback (following visuomotor rotation training). HC had significantly larger angular deviations for all target locations in the testing phase compared to baseline (15°: paired t-test t (30) = 18.3, *p* < 0.001, Cohen’s d = 3.29; 75°: paired t-test t (30) = 11.4, *p* < 0.001, Cohen’s d = 1.98; 135°: paired t-test t (30) = 10.23, p < 0.001, Cohen’s d = 1.84). For SZP, baseline and testing movements to the 135° target (baseline: –0.2 ± 0.4°, testing: 1.0 ± 0.5°) were not significantly different (paired t-test t (29) = 1.82, *p* = 0.10 Cohen’s d = 0.32). This was not the case for movements to the 15° target (paired t-test t (29) = 17.05, *p* < 0.001, Cohen’s d = 3.05) or 75° target (paired t-test t (29) = 4.03, *p* = 0.01, Cohen’s d = 0.88). (* represents *p* < 0.05, *** represents *p* < 0.001 and ns represents not significant, *p* > 0.05, for paired two-tailed t-tests).

**Figure 5.**
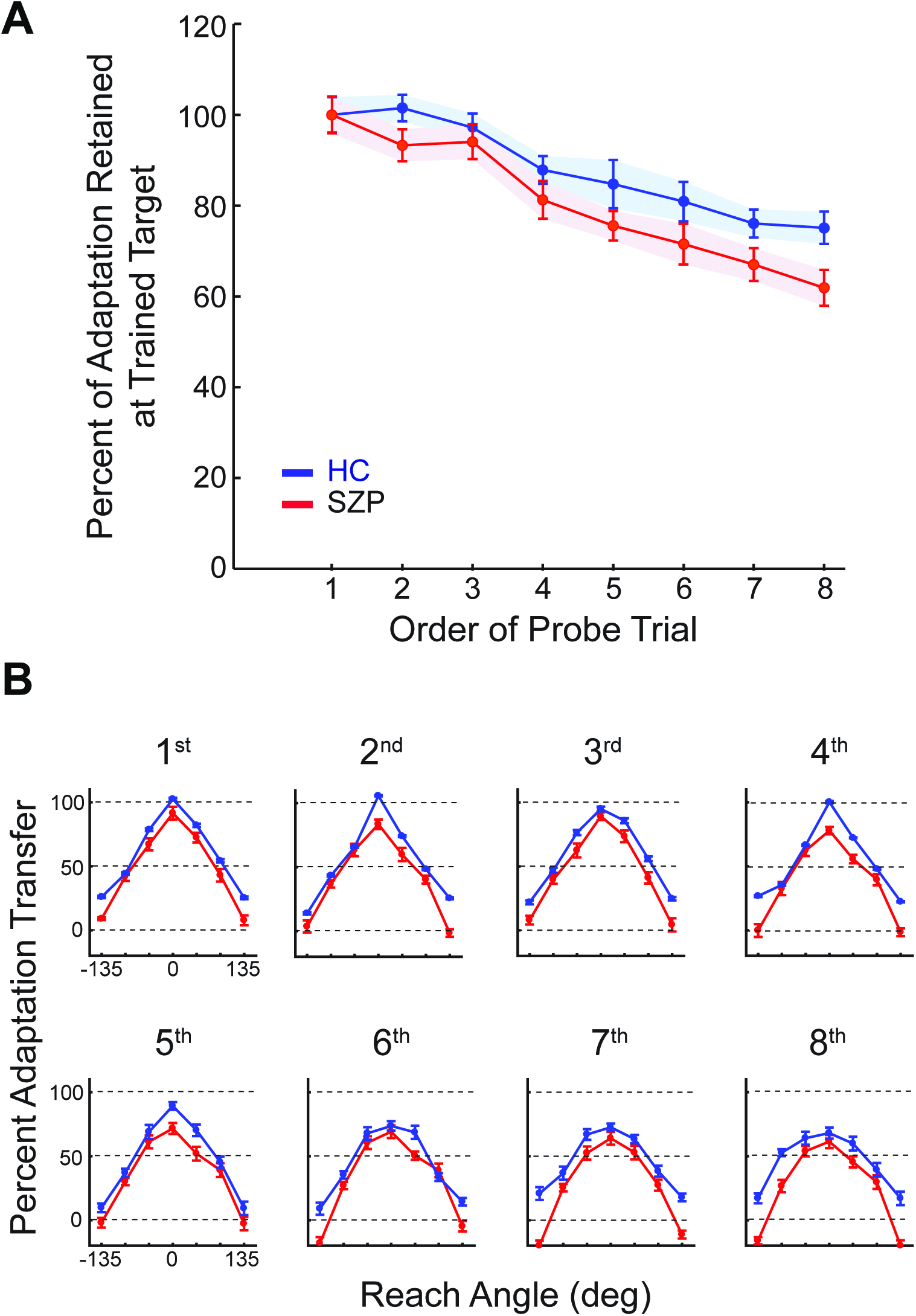
Temporal stability of spatial generalization. **(A)** Temporal decay of adaptation at the trained target was determined over generalization probe order (1 through 8). Both SZP and HC (red and blue traces, respectively) demonstrated a decrease in adaptation at the trained target over the 8 generalization probe trials. The retention of adaptation was normalized by the group mean so that on the first generalization probe trial the adaptation was 100% and the retention on the subsequent probe trials (2-8) are relative to the first probe. Filled circles represent mean values and vertical lines represent standard error. **(B)** Generalization functions over probe order. We derived the mean adaptation transfer during the testing phase for each target location, separated by order of the generalization probe trial. This resulted in eight generalization curves depicting the spatiotemporal stability of adaptation transfer over the order of the movement in the sequence.

To quantify the spatial influence on the amount of adaptation transferred, we derived a percentage of adaptation retained across spatial targets. This measure is based on the difference in the amount of adaptation transferred (during the testing phase) to a peripheral target (135°) compared to the adaptation at 0° (Figure 3A). Essentially, this was the percent of the angular deviation at 0° that was still applied at the peripheral target. Thus, a value of 100% indicates that the transfer amount was the same between the 0° and 135° target. Values between 100% and 0% indicate the amount transferred between the 0° and 135° target. Finally, a negative percentage indicates that the movements were opposite that of the compensation (i.e., the movement direction was in the same direction as the perturbation signifying no adaptation/compensation at the peripheral target. As shown in Figure 4, a number of SZP displayed a movement trajectory < 0 during testing at the 135° targets. This was never the case for HC.)

### Statistical Analyses

In order to assess group effects and differences in adaptation levels, we determined movement angles during early, middle and late adaptation, and applied a mixed-design ANOVA, including group as a between-subjects factor and trial bin (early, middle or late) as a within subjects factor in the model. We conducted a group by workspace location (left or right) by target location (±15°, ±75° and ±135°) ANOVA. Greenhouse–Grier correction was used when the sphericity assumption was not met. To examine prior associations of interest between perceptual performance measures and clinical symptoms, as well as sense of agency assessments, we evaluated correlations at a corrected: Bonferroni correction was applied to control for Type I error rate. All t-tests were two-tailed. Due to the observed group differences in WRAT (IQ estimate) scores and marginal differences in years of education between the two groups, we conducted analyses of covariance using these scores as covariates to verify our results. We report analysis of variance results because covariate analyses yielded the same results. All statistical analyses were performed using MATLAB and JASP software (JASP (Version 0.8.5); jasp-stats.org).

## RESULTS

We examined the ability of Schizophrenia patients (SZP) and healthy control subjects (HC) to adapt reaching arm movements to rotated visual feedback, and apply the learned motor recalibration to untrained movement directions. First, there was an initial baseline phase during which participants made 9 cm reaching movements (with and without visual feedback) to peripheral targets at various radial locations (0°, ±15°, ±75° and ±135° away from the trained target location, see Figure 1C). Second, in the training phase, the endpoint visual feedback of the movements made to a single target (0°) was rotated around the hand path by either (clockwise, CW) or *θ* = ‒30° (counterclockwise, CCW) (see Materials and Methods). Finally, after adaptation in the training phase, participants completed a series of generalization and retraining sequences (depicted in Figure 1D) to quantify the retention (0°) and ability to transfer the learned adaptation to the untrained target locations (±15°, ±75° and ±135°). The generalization probe trials were completed without visual feedback in order to compare these movements to the initial baseline trajectories.

### Comparison of movement metrics during baseline

Table 2 shows the baseline movement metrics that we compared between groups over the testing angles, (using a repeated measures ANOVA within each group for Feedback by Target Location) along with the accompanying statistics. We also conducted an overall Group by Visual Feedback by Target Location ANOVA for each measure. (To avoid redundancy of reporting for each measure, we omit some non-significant statistics within the text, as they are reported in detail in Table 2). Although SZP had lower peak velocities, longer movement durations, and slightly higher end-point variability, these differences were not significantly different from HC. Within each group, when comparing baseline movements with full visual feedback to those without visual feedback, we found a significant effect of Target Location (increasing radial distance away from the vertical midline) on peak velocity in both groups [HC, (F_1,29_ = 18.59, *p* < 0.001); SZP, (F_1,28_ = 5.24, *p* = 0.002)] along with a significant effect of Target Location on end-point variability in SZP (F_1,28_ = 5.50, *p* = 0.001). However, we only observed a Visual Feedback by Target Location effect for peak velocity in SZP (F_1,28_ = 6.11, *p* < 0.001). Peak movement velocity in trials with no visual feedback appeared to increase slightly with increasing radial distance away from the vertical midline, which likely resulted in increasing the end-point variability. There were no main effects of visual feedback in either group for any kinematic measure.

**Table 2.**
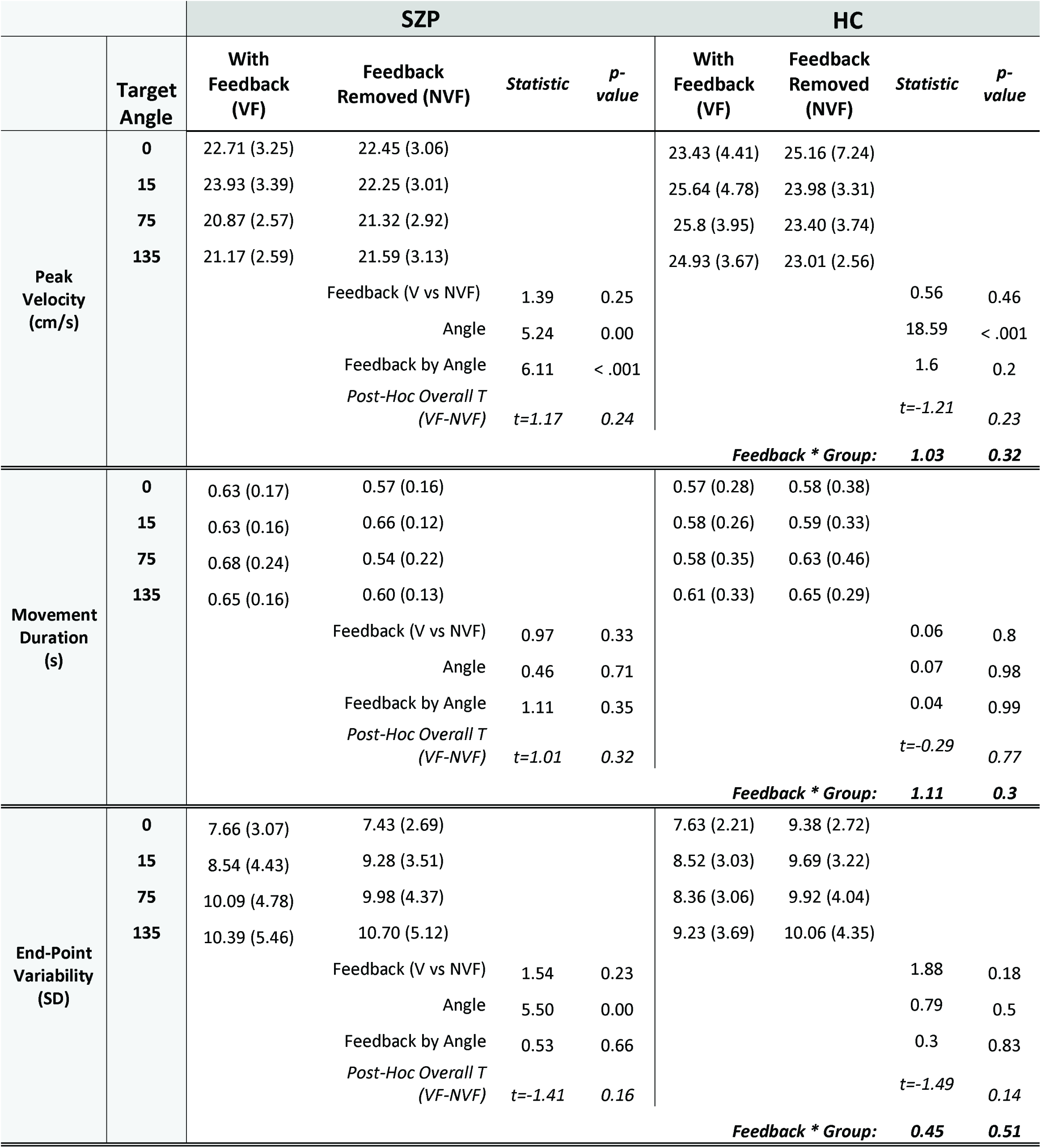
Summary and comparison of kinematic data

### The time course of visuomotor adaptation is similar for HC and SZP

In the training phase, when rotated end-point visual feedback was provided, both groups were able to compensate by adjusting their movement direction by approximately 86% of the applied perturbation amount by the end of the training (Figure 2A and B). As expected, for both groups, mean adaptation levels increased throughout the course of training (main effect of Adaptation Period on percent adaptation, (F_1,59_ = 223.13, *p* < 0.001, η^2^_p_ = 0.79). Percent adaptation to the visuomotor rotation was not significantly different between groups for early, middle, or late periods of training [Early: HC = 40.4 ± 15.5%; SZP = 32.4 ± 17.1%; Middle: HC = 83.6 ± 18.1%; SZP = 79.0 ± 20.2%; Late: HC = 86.7 ± 14.4%; SZP = 85.7 ± 14.7%]. Correspondingly, there was no significant effect of Group on the percentage of adaptation during training (F_1,59_ = 0.90, *p* = 0.34, η^2^_p_ = 0.03). Additionally, there was no interaction between Group and Adaptation Period (F_1,59_ = 1.97, *p* = 0.14, η^2^_p_ = 0.007).

### Generalization of visuomotor adaptation is narrower for SZP compared to HC

During the testing phase, in order to assess the spatial transfer of adaptation to untrained targets, we examined the amount of transfer to the seven targets, spanning –135° to 135° from the trained movement direction. Figure 3A shows the generalization curves which represent the percentage of transfer to each of the targets for each group (HC are represented by blue symbols, SZP are red). We first assessed whether there was any asymmetry in generalization with regards to the workspace location. A Group by Target Location (distance away from trained direction) by Workspace Location (left side versus right side of the workspace) ANOVA revealed a significant main effect of Target Location (F_1,59_ = 140.77, *p* < 0.001, η^2^_p_ = 0.71), Group (F_1,59_ = 65.46, *p* < 0.001, η^2^_p_ = 0.52), and a Group by Target Location effect (F_1,59_ = 3.81, *p* = 0.03, η^2^_p_ = 0.13), but no significant asymmetry in Workspace Location (F_1,59_ = 1.89, *p* = 0.18, η^2^_p_ = 0.03), and no Group by Workspace Location by Target Location interaction (F_1,59_ = 0.002, *p* = 0.99, η^2^_p_ = 0). Thus, the target location (distance from the trained direction), not workspace location (left or right of trained target location), significantly influenced the amount of spatial generalization.

Overall, for both groups, the amount of generalization decreased for targets further away from the trained target location, and this decrease was significantly greater for SZP compared to HC. In order to assess the spatial changes in adaptation transfer during the testing phase, we derived a metric to quantify the relative changes in generalization. First, due to the lack of a significant difference in the percent transfer across workspace locations, we grouped the adaptation transfer across locations in order to quantify the transfer relationships over the absolute distances from the trained target direction. The generalization data were then normalized so that at the trained direction (0° target location) the adaptation was 100% (see Materials and Methods), and the adaptation to targets at 15°,75° and 135° away from the trained direction are relative to that at 0°. Briefly, we normalized each subject’s adaptation at 0° by the group mean, and the transfer to other targets relative to this mean value. Thus, we normalized the transfer based on the adaptation at the trained target (0°) in order to cancel any differences in the trained recalibration between groups. The curves in Figure 3B show this normalized percent transfer. A Group by Target Location analysis revealed significant effects of Group (F_1,59_ = 19.53, *p* < 0.001, η^2^_p_ = 0.25), Target Location (F_1,59_ = 238.32, *p* < 0.001, η^2^_p_ = 0.80), and Group by Target Location interaction (F_1,59_ = 8.13, *p* < 0.001, η^2^_p_ = 0.13). The amount of normalized generalization was significantly different between the groups at the 75° target [(HC = 44.3 ± 3.8%, SZP = 32.9 ± 5.4%) t(59) = 2.93, *p* = 0.048, Cohen’s d = 0.64] and at the 135° target [(HC = 27.9 ± 2.3%, SZP = −4.1 ± 2.2%); t(59) = 10.10, *p* < 0.001, Cohen’s d = 2.57]. The amount of normalized transfer at 15° was lower for SZP, but this was not statistically significant [(HC = 72.8 ± 3.1%, SZP= 68.2 ± 4.8%); t(59) = 0.93, *p* = 0.21, Cohen’s d = 0.31].

Considering the group differences within the testing phase, we also examined differences between the baseline phase movements and testing phase (post-adaptation) movements in both groups (Figure 4) for target locations away from the trained direction (15 °,75° and 135°). A Group by Target location by Phase (Baseline and Testing) ANOVA revealed main effects of Target Location (F_1,59_ = 103.9, p < 0.001, η^2^p = 0.63), Phase (F_1,59_ = 592.37, p < 0.001, η^2^p = 0.91), and interaction effects of Phase by Group (F_1,59_ = 63.03, p < 0.001, η^2^p = 0.51) and Target Location by Phase (F_1,59_ = 111.55, p <0.001, η^2^p = 0.65). The Target Location by Group interaction (F_1,59_ = 2.84, p =0.06, η^2^p = 0.05) and the three-way Target Location by Phase by Group (F_1,59_ = 0.03, p = 0.98, η^2^p < 0.001) were not significant. Importantly, HC had significantly larger angular deviations post-adaptation for all target locations (Main effect of Group (F_1,59_ = 21.66, p < 0.001, η^2^p = 0.27). Collectively, these measures of spatial generalization demonstrate that SZP had a narrow spatial range over which the motor recalibration was applied, with movements to the furthest targets (135°) not significantly different from baseline movement (See Figure 4)

### Decreased spatial generalization in SZP is not due to a faster decay in the adaptation

Zhou and colleagues (2017) found in healthy subjects that generalization around the trained direction (± 15°) significantly decreased with the delay between movements and distance from the trained target, while locations further away displayed near constant spatiotemporal transfer. In order to quantify this decrease, we first obtained measures of temporal decay of adaptation at the trained target location (0°) over the order of generalization probe trials (1 through 8 in the testing phase). Similar to the normalized percent transfer of adaptation in Figure 3B, we normalized the adaptation values so that on the first generalization probe trial the adaptation was 100% (see Materials and Methods), and the adaptation on the subsequent probe trials (2-8) are relative to the first. In other words, we normalized each subject’s adaptation on the first probe trial by the group mean, and the amount on the subsequent probes relative to this mean value. Thus, this normalization canceled any differences in the trained recalibration between groups on the first probe trial in order to compare the rate of decay. As depicted in Figure 5A, we observe that the retention in adaptation at the trained target decreased for both SZP and HC over the eight probe trials. To examine these relationships, we conducted a Group by Probe Order analysis. There were main effects of Probe Order (F_1,59_ = 41.45, *p* < 0.001, η^2^_p_ = 0.45), but not Group (F_1,59_ = 3.93, *p* = 0.05, η^2^_p_ = 0.08) or Group and Probe Order (F_1,59_ = 0.69, *p* = 0.63, η^2^_p_ = 0.01). Thus, although adaptation levels for SZP were lower, there was no significant difference between groups in the retention of the motor recalibration over the successive generalization probe trials.

To further assess whether HC and SZP differed in spatial generalization of adaptation over time/consecutive movements, we derived mean values for movement angles (during the testing phase) for each target location, separated by order of trial presentation (there were eight consecutive generalization probe trials in the testing phase, see Materials and Methods). This resulted in eight generalization curves (displayed in Figure 5B), showing the temporal stability of adaptation transfer over order of presentation. There was a clear decrease in the overall generalization patterns for both groups over the consecutive trials. Consistent with the results in Figure 3A, the generalization curves for the HC (blue traces) were always above the respective SZP curves (red traces). This suggests that the difference in generalization between HC and SZP over all probe trials in Figure 3 was consistent across time/consecutive movements. That is, any temporal difference in the retention of adaptation at the trained target (Figure 5A) could not account for variations in the pattern of spatial generalization (Figure 3). This was supported by a Group by Target Location (collapsed over left and right workspace for 15°, 75° and 135°) by Probe Order ANOVA. There were significant main effects of Probe Order (with Greenhouse-Geisser correction, F_1,59_ = 7.26, *p* < 0.001, η^2^_p_ = 0.11), Target Location (with Greenhouse-Geisser correction, F_1,59_ = 219.77, *p* < 0.001, η^2^_p_ = 0.78) and Group (F_1,59_ = 5.59, *p* < 0.021, η^2^_p_ = 0.08). However, all interactions were not significant (each with Greenhouse-Geisser correction): Probe Order and Target Location (F_1,59_ = 7.26, *p* = 0.08, η^2^_p_ = 0.03), Probe Order and Group (F_1,59_ = 1.30, *p* = 0.26, η^2^_p_ = 0.03), Target Location and Group (F_1,59_ = 2.31, *p* = 0.09, η^2^_p_ = 0.04), and Probe Order, Target Location and Group (F_1,59_ = 0.37, *p* = 0.96, η^2^_p_ = 0.01). Thus, differences between the HC and SZP groups in the spatial transfer of adaptation were not due to any persistent temporal difference in the retention of adaptation.

### Correlations with clinical SZ symptoms and subjective traits

For each subject we determined the extent of adaptation transfer (the percentage of adaptation retained across spatial targets, see Materials and Methods) and examined associations of this measure with clinical symptoms and SoA measures. For the SZP group, a critical p value of 0.008, equivalent to a Bonferroni correction of p < 0.05 for 6 pair-wise tests was set [One generalization measure (Spatial transfer between the 0° and 135° target), three SoA measures (physical, mental and Total SoA) and three symptom measures (positive, negative and a grouped total psychotic symptoms)]. We found strong associations between the amount of spatial change in adaptation transfer (the percentage of adaptation retained from 0° to 135°) and total positive symptoms as measured by the PANSS (*r* = −0.52, *p* = 0.003, Figure 6A). The plot indicates that SZP presenting with more severe positive symptoms (larger scores) had greater difficulty in the ability to transfer the learned visuomotor recalibration to untrained targets, as indicated by the negative percentage of adaptation retained. Notably, spatial changes in adaptation transfer were also associated with a grouped total psychotic symptoms score that included delusions, hallucinations, unusual thought, disturbance of volition and grandiosity) (*r* = −0.51, *p* = 0.004).

**Figure 6.**
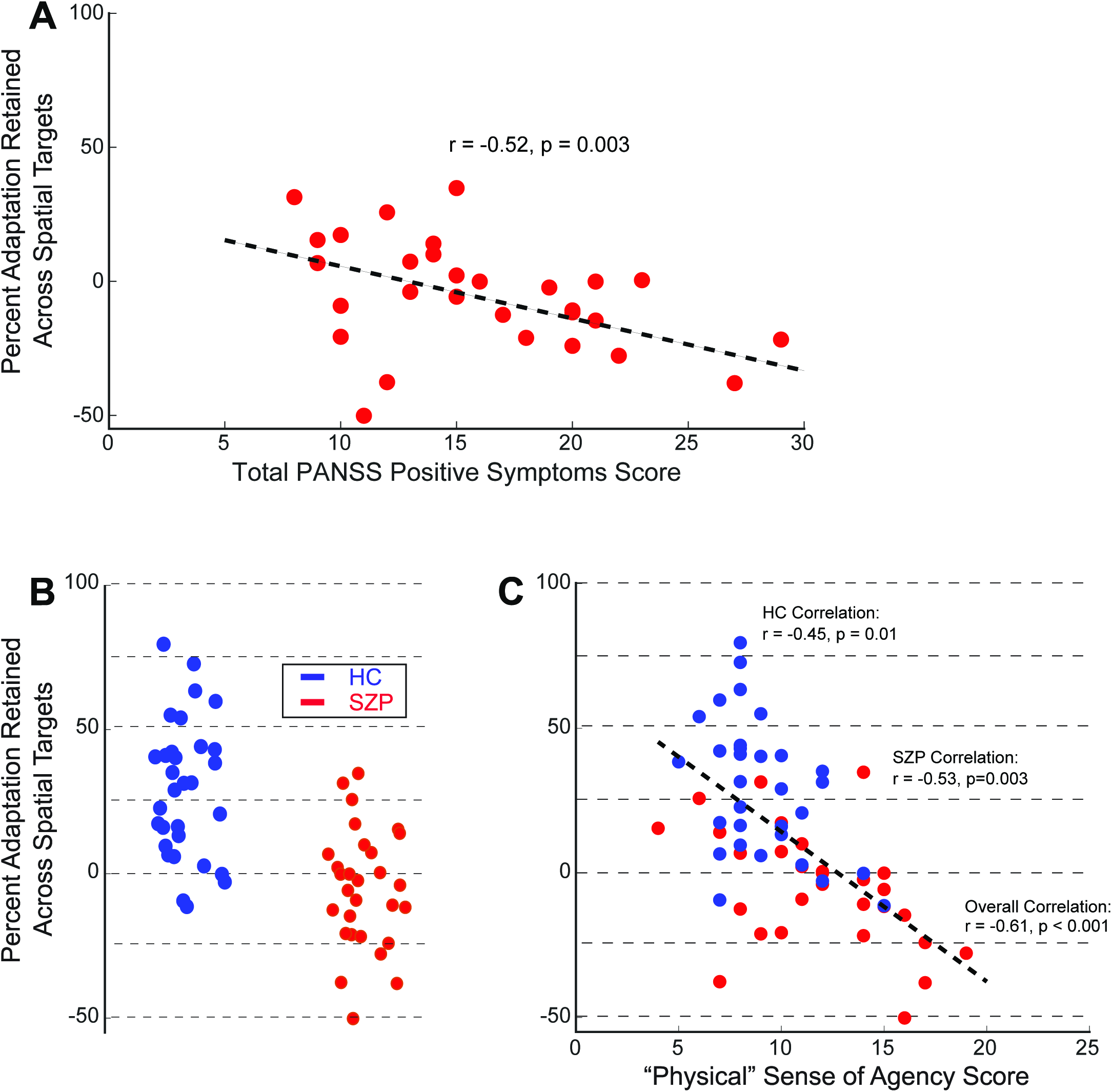
Associations between the spatial changes in adaptation transfer, PANSS positive symptoms and physical SoA. We derived a measure, *percent adaptation retained across spatial targets*, based on the angular deviation at the 0° target and the 135° target. A value of 100% indicates that the transfer amount was the same for the two targets, while a value between 100% and 0% indicates the transfer amount from 0° to 135°. **(A)** The scatter plot shows the relationship between this measure of adaptation transfer and Total PANSS positive symptoms for SZP, indicating that SZP presenting with more severe positive symptoms (larger scores) had greater difficulty in the ability to transfer the learned visuomotor recalibration to untrained targets. Each filled red circle represents a single SZP and the black dashed line represents the linear regression between the two measures. **(B)** The plot shows the overlap in the spatial change in adaptation transfer between the HCs (filled blue circles) and SZPs (filled red circles). The changes in transfer for some HC were similar to that of the SZP group, and vice versa. **(C)** The scatter plot shows the relationship between the spatial change in adaptation transfer as a function of the ‘Physical’ sense of agency measure. The black dashed line is the linear regression between these two measures for the entire sample (SZPs and HCs combined). The Spearman r and p values are displayed for the respective groups and over the entire sample.

We did not find any association between the spatial transfer measure and negative symptoms. In addition to the strong correlation to total positive symptoms, the spatial transfer of adaptation correlated (*r* = −0.53, *p* = 0.003) with the measure of ‘Physical’ sense of agency for SZPs (Sense of Agency involving somatic experiences, see Materials and Methods) (Figure 6C).

As shown in Figure 3, there was variability in the adaptation transfer within both the HC and SZP groups. Subsequently, there was a modest amount of overlap in the spatial transfer of adaptation across the two groups (Figure 6B). That is, there were some HC subjects that demonstrated a difficulty in generalization similar to the SZP group, and vice versa. Surprisingly, within the HC group (critical p value of 0.02, equivalent to a Bonferroni correction of p < 0.05 for 3 pair-wise tests (Total SoA, Physical and Mental SoA and one generalization measure), we also found an association between the percent change in the spatial adaptation transfer and the measure of ‘Physical’ sense of agency (*r* = −0.45, *p =* 0.01). There was a strong correlation across both HC and SZP, (*r* = −0.61, *p* < 0.001) suggesting a continuum in motor adaptation transfer and sense of agency across the two groups.

## DISCUSSION

We examined the ability of Schizophrenia patients (SZP) and healthy control (HC) subjects to spatially generalize the adaptation of reaching arm movements to different directions. Based on well-documented defects in the internal monitoring of movement, we hypothesized that SZP would recalibrate movement based on the visual feedback, but show an impairment in the ability to transfer this recalibration to novel conditions when feedback was absent. We found that compared to HC, the motor recalibration in response to visual feedback perturbations (a rotation with respect to the actual movement vector) was similar for SZP. However, SZP were not able to generalize the motor adaptation to the same extent as HC. This difference was not due to deficit in the retention of adaptation at the trained target; the level of transfer to untrained targets over consecutive probe trials was always less for SZP compared to HC. Importantly, our measure of motor adaptation generalization was correlated with positive symptoms such as delusions and hallucinations in SZP, and a measure of ‘physical’ sense of agency (SoA) in both SZP and HC, suggesting that the impaired transfer of motor learning was related to an altered sense of action ownership. Our results provide insight into how abnormal ownership of thoughts and actions adversely affect how humans model, interpret and interact with the environment.

### Sense of agency and learning mechanisms during visuomotor adaptation

Schneider (1920) noted almost a century ago that SZP exhibit deficits in action attribution. It has been postulated that action attribution and SoA depend on sensory prediction that compares predictions made by an internal model of the motor system and sensory signals resulting from the corresponding action (Frith *et al*. 2000; Georgieff and Jeannerod, 1998; Sperry, 1950; von Holst and Mittelstaedt, 1950): a correspondence between predictions and sensory signals leads to attribution of the observed effects to an internal source and a mismatch leads to an external designation. Motor adaptation is a form of learning that involves accurate action attribution; the movement perturbation can induce a sensory prediction error (the perceived difference between the expected and actual movement consequences), which can be used to update the internal model prediction and adjust subsequent motor commands (Shadmehr *et al*., 2010). Adaptation to visuomotor rotation at least partially depends on the integration of both internal prediction and external sensory feedback. It follows that generalization of the learning to new locations without visual feedback would depend in part on internal prediction, with imprecise prediction associated with a disturbed sense of agency. In our study, SZP and HC demonstrated similar adaptation during the training phase, in which feedback was present (Figure 2). This suggests that the sensory feedback component of the predictive mechanism in SZ was largely intact and sensitive to the feedback perturbation. However, despite similar adaptation, SZP showed negligible generalization to untrained targets when feedback was removed, whereas HC applied approximately 25% of the learning to the farthest untrained targets (135°). Consistent with at least a partial role of internal model prediction in the transfer of motor learning, we observed that deficits in generalization in SZP were associated with the severity of psychosis and an abnormal sense of agency. Importantly, HC who also showed less generalization also had a reduced sense of physical agency.

Based on the aforementioned deficits in sensory prediction error (Bansal *et al*. 2018b; Daprati *et al*. 1997; Lindner *et al*. 2005; Martinelli *et al*. 2017; Shergill *et al*. 2005, 2014) it would be natural to ascribe abnormal generalization in SZP to deficits in the utilization of sensory prediction errors. However recent evidence suggests a different interpretation. Both implicit and explicit processes have been shown to contribute to visuomotor adaptation and generalization of learning (Day *et al*. 2016; Hegele and Heuer, 2010a, b; Mazzoni and Krakauer, 2006). For example, for recalibration based on altered visual feedback, Taylor and colleagues (2014) showed that overall adaptation is a combination of (1) explicit mechanisms, characterized by intentional control of movement aiming direction and (2) implicit mechanisms, characterized by corrections based on sensory prediction errors. Additionally, McDougle *et al*. (2017) found that the pattern of transfer of adaptation across the workspace for HC is different for the various learning mechanisms; implicit learning generalizes in a Gaussian manner, while explicit learning is transferred in an approximately uniform manner over the workspace (see also Heuer and Hegele, 2011). Interestingly, the authors found that generalization to far targets (> 67.5≥ from the trained target direction) was largely driven by the explicit component while generalization to nearby targets (similar to the 15° target in the current study) was driven by both implicit and explicit components of learning.

Based on these studies, one interpretation of the generalization deficit in SZP is a reduced ability to utilize explicit or plan-based generalization to new targets. However, there are issues with this interpretation. First, it should be noted that the difference in generalization between HC and SZP was not uniform (Figure 3B); the separation in transfer between the two groups increased with distance away from the trained target direction, suggesting that the deficit in SZP is more complex than simply a reduction in a purported uniform explicit learning component. Second, it is possible that the combination of explicit and implicit learning mechanisms across the workspace is different in SZP compared to HC, given the known deficits in planning and intentional control in SZP (e.g. Frith *et al*., 2000; Voss *et al*., 2010, 2017). Thus, patients may have relied on less effective implicit processes (sensory prediction) in applying the recalibration to peripheral targets. A follow-up study is necessary to directly separate different learning mechanisms and their drivers (Izawa and Shadmehr, 2011; Taylor *et al*. 2014; McDougle *et al*. 2017) and provide further insight into the specific motor learning deficits and subsequent generalization patterns we have presented.

### Adaptation mechanisms and neural substrates

In general, implicit learning during motor adaptation is facilitated through sensory prediction mechanisms involving in the cerebellum (Crapse and Sommer, 2008; Houk *et al*. 1996; Kawato, 1999; Wolpert *et al*. 1998) that modify neuronal activity in the posterior parietal and motor cortices to reduce the sensory prediction error (Tanaka *et al*. 2009). Evidence from various neurophysiological, clinical and behavioral studies highlight the roles of the cerebellum, parietal and motor cortices during various phases of motor adaptation to perturbed visual feedback (e.g., Della-Maggiore and McIntosh, 2005; Galea *et al*. 2011; Hadipour-Niktarash *et al*. 2007; Krakauer *et al*. 2004; Paz *et al*. 2003; Tseng *et al*. 2007; Yavari *et al*. 2016). For example, patients with cerebellar lesions show a pronounced impairment in their ability to adapt to novel perturbations (e.g., Donchin *et al*. 2012; Izawa *et al*. 2012; Rabe *et al*. 2009; Smith and Shadmehr, 2005).

The parietal lobe and cerebellum are involved in various aspects of sensorimotor prediction such as recognizing the sensory consequences of action and distinguishing self-produced and external movements, which are functions related to SoA (Blakemore and Sirigu, 2003). Imaging studies have implicated several brain areas associated with SoA (Blakemore *et al*. 2001; David *et al*. 2008; Farrer and Frith, 2002; Farrer *et al*. 2003; Fink *et al*. 1999; Jeannerod, 2004; Leube *et al*. 2003), including the ventral premotor cortex, supplementary motor area and cerebellum, which also constitute a network of sensorimotor transformations and motor control. Evidence from the tactile (Blakemore *et al*. 1998, 1999, 2001), visuomotor (Synofzik *et al*. 2009) and force processing domains (Martinelli *et al*. 2017; Shergill *et al*. 2005, 2014) indicate that efficient utilization of sensory prediction is crucial to maneuver within the environment. When sensory prediction fails, as seen in neuropsychiatric disorders like SZ, disturbances in SoA, action inference and modulation arise. Impaired predictive signals emanating from the cerebellum could contribute to this failure, thus driving the use of alternative learning mechanisms.

In SZ, a disrupted prefronto-thalamo-cerebellar circuit (Andreasen *et al*. 1998; Barch, 2014; Giraldo-Chica *et al*. 2018) has been proposed to play a role in the pathophysiology. There is evidence for reduced cerebro-cerebellar connectivity in higher level association networks (e.g. ventral attention, salience, frontoparietal control system) and increased cerebro-cerebellar connectivity in somatomotor and default mode networks (e.g., self-referential and undirected spontaneous, mental activity, Buckner *et al*. 2008; Gusnard *et al*. 2001). It is reasonable that alterations between these networks could give rise to disturbances in SoA as related to somatosensory and motor control predictive functions. It has been postulated that the aberrations in somatosensory and motor control function in SZ may stem from cerebellar dysfunction or disconnection (Andreasen *et al*. 1998; Schmahmann and Sherman, 1998). Overall, the underlying mechanisms of sensory prediction error mechanisms appear to be aligned with this dysconnection framework (Friston *et al*. 2016).

Here, we provide support for these theories by demonstrating a strong link between deficits in the generalization of visuomotor adaptation, and aberrant SoA, both of which are thought to partially involve the cerebellum, and may be affected by cerebellar circuit abnormalities. Future studies in non-psychiatric and clinical populations are needed to clarify the interplay of sensorimotor function as related to action, and the functional basis of SoA as related to other cognitive processes. Additionally, future work is needed to examine the spatiotemporal properties of different adaptation mechanisms and differential effects on generalization patterns. This is of clinical relevance as central processes of sensorimotor integration are crucial for correct attribution of agency and ownership to movements and actions.

## ACKNOWLEDGEMENTS AND DISCLOSURES

We are grateful to the veterans at the Washington DC VA Medical Center who participated in this study. The authors report no financial interests or potential conflicts of interest. SB is currently affiliated with the University of Maryland School of Medicine, Maryland Psychiatric Research Center, Catonsville, MD.

